## Supplementary Information (SI) for "Substrate evaporation drives collective construction in termites"

#### S.I Experimental specimens

**Termite species** In our experiments, we studied the building behavior of termites *Coptotermes gestroi*. *Coptotermes gestroi* is a soil-nesting wood-feeding termite originating from South-East Asia [*S10*]. However, as a result of the increase of global human activity, it has spread in Asia and to other parts of the world including Africa, Europe, North America, Central America and South America [*S14*]. As an exotic pest it is primarily found in populated urban environments, where it feeds on human-made structures [*S13*]. In its natural habitat C. gestroi feed on dead trees and wooden debris on the soil surface and build their nest underground, although "aerial" infestations have also been recorded in human made structures, where the nest have no contact with the ground. Whether the nest is aerial or subterranean, the internal nest structure is similar to the nests of other termites in the *Coptotermes* genus, and comprises a "scaffold" of interconnected pillars (see figure S1). This structure is the result of a building process, as evidenced by the fact that its material composition is different from the composition of the surrounding soil and comprises some stercoral carton. Colonies of *C. gestroi* are estimated to be within the range of 100,000 to 4,000,000 individuals [*S1*].

**Captive colonies** For most experiments, termites were collected from the same master captive colony (c22) of *Coptotermes gestroi* hosted at the LEEC laboratory (Villetaneuse, France) in a tropical room with constant temperature (26±2°C) and

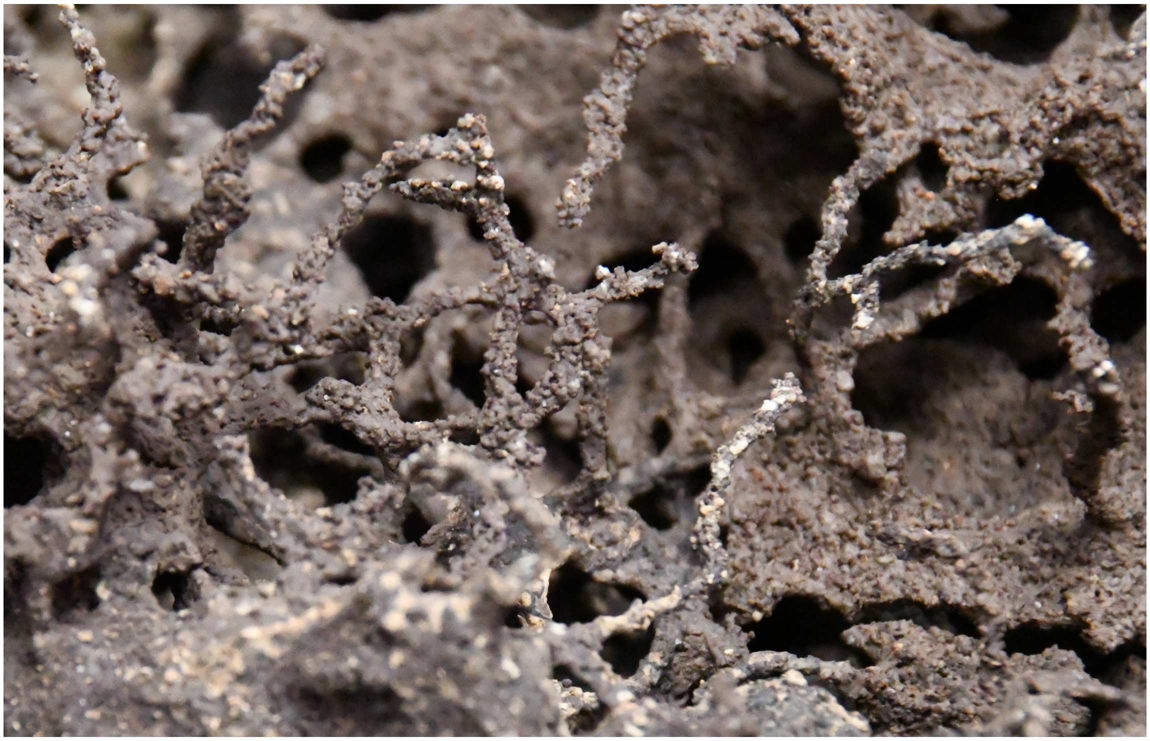

Figure S1: Close-up on pillar structures built by our captive colony of *Coptotermes gestroi* within the plastic barrel that hosts the full colony.

relative humidity (70±10%). Only experiments 57, 90, 91, 92, 93 and 94 were run using termites from a second master colony (c21). Experiments were performed in the same tropical room in which the master colonies were housed to reduce the environmental stress on the experimental populations. Workers were attracted with humid towels and gently shoveled with a pencil on a plastic tray. Groups of 50 workers and 5 soldiers were then formed using insect forceps and added to an experimental setup. Soldiers were mostly inactive during all our experiments but were included to maintain the same proportion as in a real colony (1 soldier for every 10 workers). This choice was made to limit as much as possible the factors that might affect building behavior for not being in a natural situation. Finally when running batteries of multiple experiments, distinct groups of 50 workers were formed by rounds of ten individuals taken individually from the plastic tray. This choice was made to avoid introducing any unwanted bias in the groups, coming from the fact that inactive or larger termites are usually picked first from the tray because they are easier to pick.

#### S.II Experiments summary

We ran a total of 32 experiments: 16 with pillars as topographic cues, 11 with walls, and 5 with no cues. In table S1, we report all the experiments with a few qualitative descriptors concerning the experiment outcomes. In agreement with the results shown in the manuscript, one sees that topographic cues as well as the edges of the clay disk attract depositions in a vast majority of experiments and depositions elsewhere are generally scarce or absent. The rightmost column indicates if the experiment show the appearance of spontaneous digging activity before 24 hours or before all sparse pellets have been collected. Adding sparse pellets did not always prevent termites from digging, but this method was effective in inhibiting digging activity in more than half of the experiments. Also we have highlighted in green the wall experiments where depositions focused at lateral tips of the top wall edge, an outcome that points toward surface curvature as the dominant cue compared to elevation. Note that experiments IDs in table S1 run from 48 to 94 with several numbers missing in-between. All missing IDs refer to one of the three cases: (i) preliminary experiments where the final protocol had not been adopted yet, (ii) experiments intended for the production of pellets which exhibit a “natural” distribution of size and (iii) experiments that were prematurely aborted due to problems with the recording device or the hydrating system. A visual confirmation of table S1 is provided in figures S2, S3 and S4 where we reported a snapshot of each experiment. For most experiments, pictures were taken after 24h or more and the picture time is superimposed in cyan at the top right of each snapshot.

Experiments were performed during three weeks between February and March 2021 at the LEEC laboratory in Villetaneuse. To assess the experiment variability independently of the colony activity at different times, we run batteries of up to 6 experiments. Each replica in a battery was prepared following meticulously the same protocol both in the preparation of the arena and in the collection of the specimen. Such batteries are grouped together in table S1 and labeled with the same color. One can see that variability can be high even within the same experimental battery as confirmed by qualitative descriptors and experimental snapshots.

Table S1: Summary table of experiments. Colors label in the leftmost column denote batteries of identical simultaneous experiments. Green highlighting denotes wall experiments where deposits focused at the lateral tips of the top wall edge. ID labeled with a * refer to cases where: the clay disk dried prematurely (57,74,75); hydrating holes were drilled on one half of the disk (76); a larger number of workers (n=150) was introduced in the arena.

| Exp ID | Cue type | Cues Deposits (any time) | Edge Deposits | Elsewhere Deposits | Spontaneous holes |
|---|---|---|---|---|---|
| 48 | pillars | Yes | No | Minor | 1 |
| 52 | pillars | Yes | No | No | No |
| 53 | pillars | Yes | Yes | Some | No |
| 57* | pillars | Minor | No | Some | Yes |
| 58 | pillars | Yes | Yes | Minor | No |
| 59 | pillars | Some | No | minor | Yes |
| 63 | pillars | Yes | Yes | No | No |
| 64 | pillars | Yes | Yes | No | 1 |
| 65 | pillars | Yes | Yes | No | 1 |
| 66 | pillars | Yes | Yes | No | No |
| 67 | pillars | Yes | Yes | Yes | Yes |
| 72 | pillars | Yes | Yes | No | No |
| 73 | pillars | Yes | Yes | Some | Yes |
| 74* | pillars | Yes | Yes | Yes | Yes |
| 75* | pillars | Yes | No | Yes | Yes |
| 76* | pillars | Yes | No | No | No |
| 49* | wall | Yes | No | No | n/a |
| 71 | wall | Some | Yes | Minor | No |
| 77 | wall | Yes | Yes | Some | Yes |
| 78 | wall | Yes | Yes | No | No |
| 79 | wall | Yes | Yes | Minor | No |
| 81 | wall | Yes | Yes | Some | Yes |
| 90 | wall | No | No | Yes | Yes |
| 91 | wall | Yes | Yes | No | No |
| 92 | wall | Yes | Yes | Minor | Yes |
| 93 | wall | Yes | Yes | No | 1? |
| 94 | wall | Yes | Yes | Minor | Yes |
| 83 | None | Spontaneous pillars | Yes | Minor | Non |
| 84 | None | n/a | Yes | Minor | Yes |
| 85 | None | n/A | Yes | No | No |
| 86 | None | Spontaneous pillars | Yes | No | No |
| 87 | None | n/A | Yes | Some | Yes |

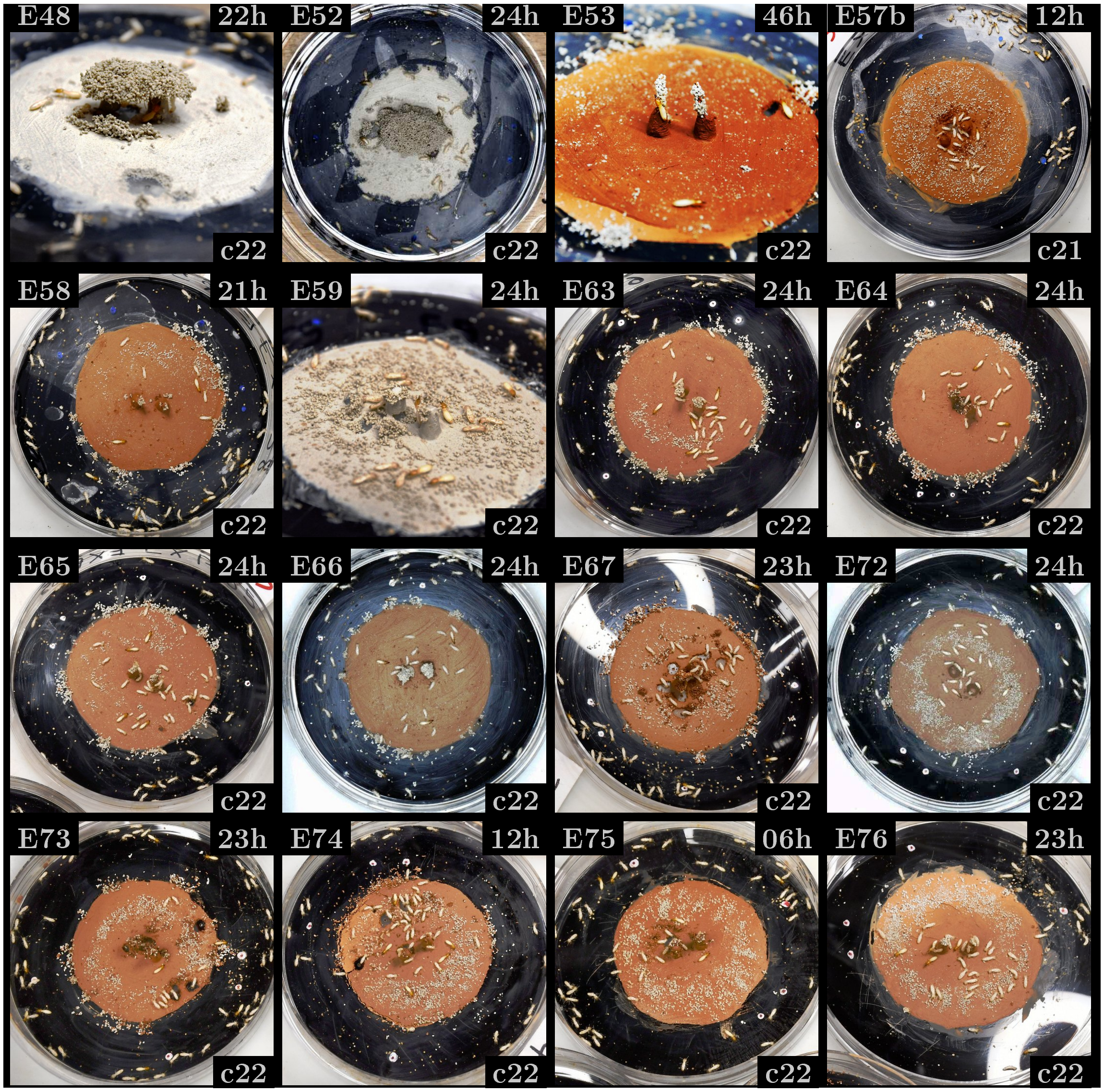


Figure S2: Snapshots of experiments with pillar cues. Superimposed labels denote: the experiment ID (top left), the time after the start of the experiment at which the picture was taken (top right) and the ID of the master colony (bottom right).

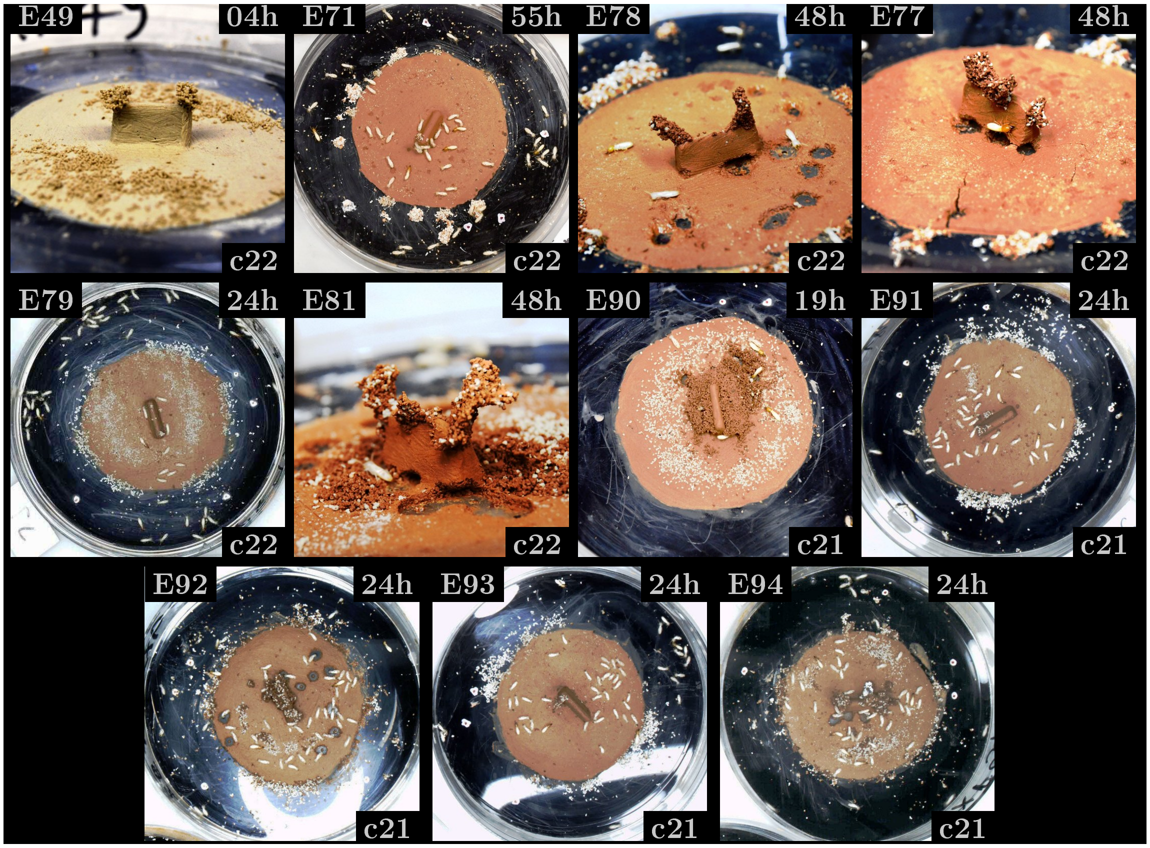


Figure S3: Snapshots of experiments with wall cues. Superimposed labels denote: the experiment ID (top left), the time after the start of the experiment at which the picture was taken (top right) and the ID of the master colony (bottom right).

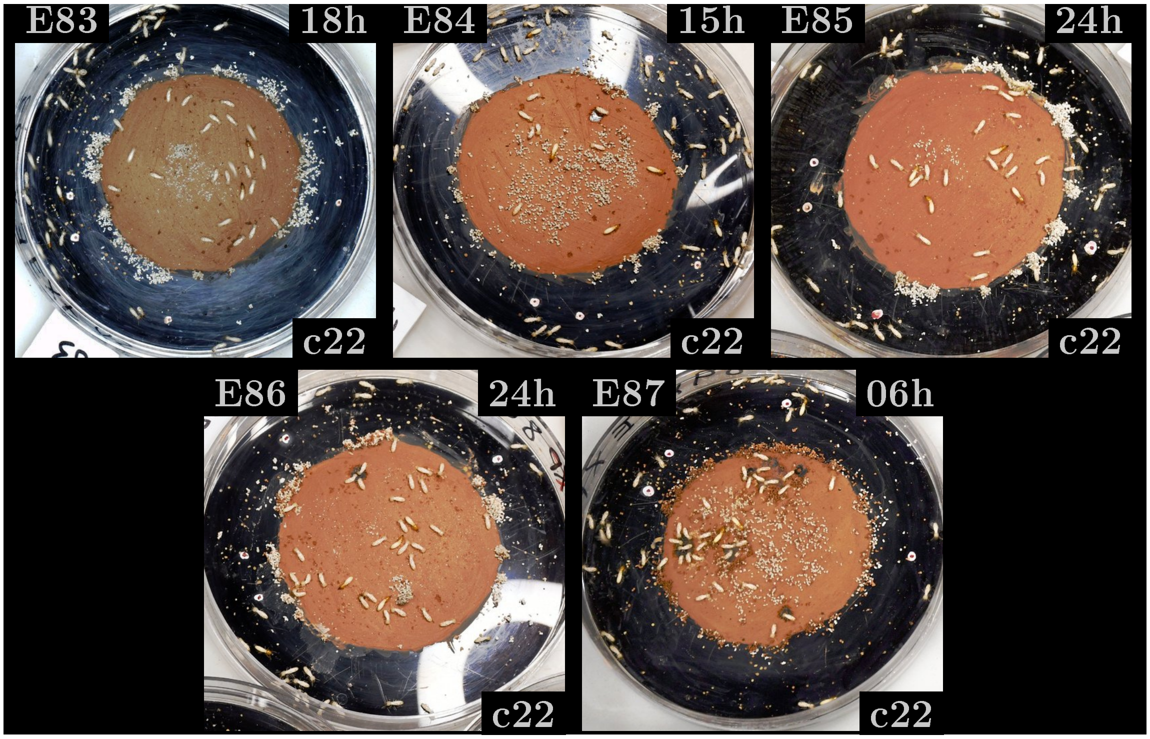


Figure S4: Snapshots of experiments with no cues. Superimposed labels denote: the experiment ID (top left), the time after the start of the experiment at which the picture was taken (top right) and the ID of the master colony (bottom right).

### S.III Image recording and analysis

A led lamp constantly illuminated the setup from above. Two types of image recording methods were used during at least 24 hours: a time-lapse at intervals of 20 seconds and a continuous video recording at 7 fps. Time-lapse were performed using a Nikon D7500 20Mp reflex camera and allowed to monitor the building behavior in the batteries of several identical experiments. Conversely, video recording was used to analyse termite motion at a finer time resolution in a single experiment of the battery. Videos were obtained using a usb-camera (MER2-1220-32U3C) with a 12 Mp 1/1.7" CMOS sensor on which we mounted a 12 mm lens (72 mm equivalent for 35 mm SLR camera).

#### Collection and deposition heatmaps

We developed an image processing pipeline that allows tracking the displacement of pellets in the arena, i.e. quantifying the collection and deposition of material. First, we consider the time-lapse or video recording and extract a sequence of images at intervals of 80 sec, which is a rough estimation of the time of a search-collection-deposition sequence. Images are then converted to greyscale and the image of the initial setup (without termites) is subtracted from all of them. At the next step, we replace each image in the sequence by the median across 10 consecutive images, i.e. each pixel takes the median value across 10 consecutive images. This operation is meant to take rid of termites that are constantly moving while pellets are usually moved only once, from the collection site to the deposition site. Notice that pellets are light grey while the arena is brown, so after subtracting the initial image, any collected pellet leaves a dark (low luminosity) trace in the sequence, while any dropped pellet leaves a bright (high luminosity) trace in the image sequence. Subsequently, we average the sequence across intervals of 10 images, thus integrating termite activity in windows of 10*80 = 800 secs.

A demo of a typical image sequence at this stage is available at this link. To quantify the amount of collected pellets, we then binarise each image at low threshold and isolate collected pellets as black regions so that they can be labeled with a connected components algorithm. We exclude components whose area is below 10 px (i.e. half the area covered by the smallest pellets) or above 400 px. The lower and upper thresholds was meant to get rid of noise and clusters of inactive termites, respectively. Finally, we convert again binary images to greyscale images, and rescale each image so that one pixel is 0.75 mm large which is the size of an average pellet. This last operation allows reducing significantly the size of our heatmap without loosing information. We have then obtained a heatmap of collection activity as a function of time. An equivalent result is obtained for the spatial distribution of depositions, simply inverting the pixel values of initial images. As a matter of fact, most of the pellets are displaced only once, thus looking at the last heatmap in the temporal sequence, one has a good estimation of the cumulative distribution of collections and depositions across the experiment: we refer to these cumulative distributions as $P(C)$ and $P(D)$ throughout the manuscript.

#### Termite occupancy

Termites motion is analyzed using the fast multi-animals tracking tool TRex [*S16*]. The ultimate aim of TRex algorithm is to identify the trajectories of each individual and record their position and velocity as a function of time. Unfortunately, TRex did not manage to preserve individual identities from our videos: the identity of single trajectories was rapidly lost and we could not rely on them to study individual termite behavior. While the identity of individual termites was not preserved, we verified that the trajectories tracked by the software truly reflect real termite locations, and as such they can be used to obtain information about the overall termite occupancy within the Petri dish. Once the trajectories are extracted by TRex, we proceed in the same way as with pellets traces, that is, we construct a heatmap of positions on the same regular grid (0.75 mm grid step) and during 800 secs intervals.

#### Conditional probabilities

To assess the randomness of collection and deposition events, we wanted to estimate the probability of these two events at a given position conditional on the probability of termite occupancy at the same position. Let us denote with $P(C|O)$ the conditional probability of collection given occupancy, then by definition:

$$P(C|O) = \frac{P(C \cap O)}{P(O)}.$$

However, a collection episode necessarily happens where a termite is, which means that collection events $C$ are a subset of occupancy events $O$, which implies:

$$P(C|O) = \frac{P(C)}{P(O)},$$

that is $P(C|O)$ is simply the ratio between collection and occupancy frequency. The same reasoning can be done for the conditional probability of deposition given occupancy $P(D|O) = P(D)/P(O)$ which is the ratio between deposition and occupancy frequency.

**Normalization of probability distribution** Note that in the plots of figure 2 (main manuscript) the frequency heatmaps ($P(C)$, $P(D)$ and $P(O)$) do not sum up to 1. Instead, we preferred to normalize the probabilities by their average value, which allows us to easily recognize high frequency areas where $P > 1$ and low frequency areas where $P < 1$. Coherently with this choice, conditional probabilities ($P(C|O)$ and $P(D|O)$) are not normalized to 1 either.

##### 3D surface scans

Before the start of each experiment, a surface scan of the setup was taken using a 3D surface scanner (NextEngine 3D Scanner ULTRA HD). This allowed initialising the numerical simulations with the same topographic cues as in the experiments, as described below in section S.VI. The scanner resolution was sufficient to characterize the curvature of topographic cues but it was insufficient to detect the edges of the clay disk. In this region an additional problem came from the fact that the Petri dish (bare) surface could not be scanned properly, likely because of the laser getting reflected two times at the top and bottom side of the plastic plate. This effect, combined with the small size of these features relative to the spatial resolution of the scanner, made it difficult to reliably measure surface curvature at the edges of the clay disk.

#### S.IV Temperature and humidity measurements

Temperature and humidity were measured using a commercial temperature-humidity probe (DHT22) connected to a Raspberry Pi micro computer. In order to not interfere with termite behavior, our temperature and humidity measurements were performed on a control experiment, which was prepared using the same protocol as the other experiments but where no termites were added. The results are reported in figure S5. The shaded area corresponds to a time interval where the probe was placed on the bottom of the Petri dish, outside the clay disk, while the rest of the time the probe was placed on the clay disk. We can observe a net decrease in temperature and an increase in humidity when moving from inside to outside the clay disk, while the overall values within the two regions do not show large variation across several hours. This is consistent with the fact that the clay disk undergoes a stationary evaporation process and the well know cooling of the substrate due to the latent heat of water.

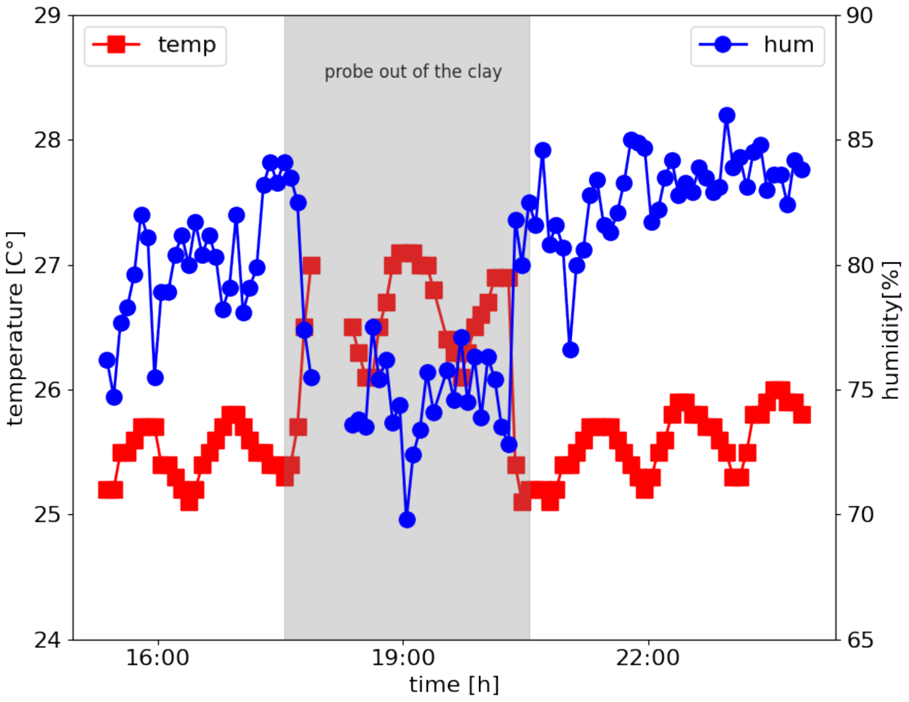


Figure S5: Temperature (blue) and humidity (red) within the Petri dish as a function of time. The shaded area refers to a time interval where the probe was placed on the bottom plate of the Petri dish but outside the clay disk. During the remaining time the probe was placed on the clay disk.

#### S.V Curvature definition

In this section, we give a mathematical definition of *surface curvature* and we explain how its sign is determined. Given a three-dimensional object, its surface can be characterized locally using the reciprocal of the radius of two osculating circles $r_1$ and $r_2$ as sketched in figure S6. One can then define the first principal curvature $k_1 = 1/r_1$ and second principal curvature $k_2 = 1/r_2$, or alternatively the mean curvature $H = (k_1 + k_2)/2$ and the Gaussian curvature $\Gamma = k_1 \cdot k_2$. Throughout the manuscript when using the term *curvature* or *surface curvature*, we always refer to the mean curvature $H$.

Provided that we have established a convention for the interior and the exterior of the surface, i.e. once the normal vector $\hat{\boldsymbol{n}}$ to the surface is drawn, principal curvatures come with a sign and so do $H$ and $\Gamma$. Indeed drawing the normal vector univocally fixes the value (and the sign) of $H$ in the form:

$$2H = -\nabla \cdot \hat{\boldsymbol{n}}.$$

Traditionally, the normal vector is oriented towards the exterior, for example thinking about a solid, the normal vector usually points toward the outer space. When this convention is adopted (which is for instance what was done in reference [*S3*]), convex objects have negative curvature and concave objects have positive curvature. Conversely, here we use the same convention as in our phase field model [*S6*] which is the opposite one: convex surfaces have positive curvature and concave surfaces have negative curvature. While, of course, experimental results and theoretical predictions do not depend on which convention is adopted, the sign of the curvature can become important when comparing results across different studies. For example, [*S3*] suggested that termite building activity is enhanced in regions of high positive

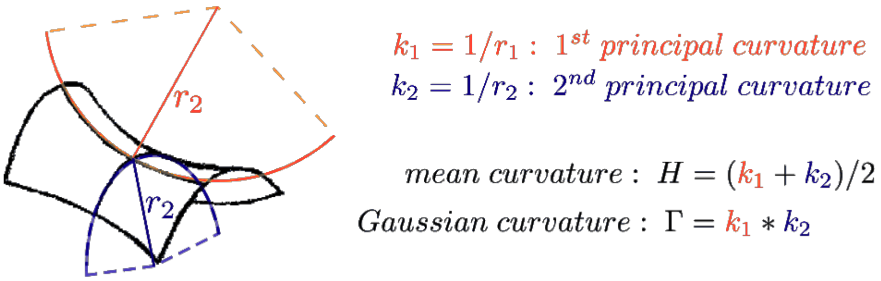


Figure S6: Sketch of curvature definition on a saddle shaped surface.

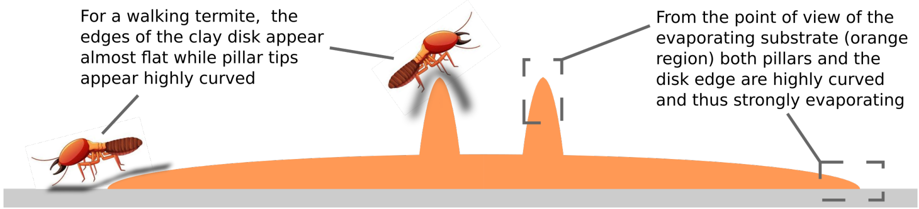


Figure S7: Sketch of the experimental setup showing the contrast between how sharp is the shape of the wet substrate (orange region) at the disk edge, and how flat the same region can appear to a walking termite.

mean curvature, that -by their sign convention- are regions of highest concavity. In the present study, we also propose that positive mean curvature attract deposition of building material, but in our case the regions of positive curvature are the most convex. Thus, the two results do not agree. However, our model predicts that while convex regions attract deposition, concave regions attract digging, and the analysis techniques used by [*S3*] do not allow distinguishing between digging and deposition actions. We then propose that what was observed in [*S3*] is a correlation between curvature and digging activity which is in agreement with our model.

### S.VI Humidity diffusive model

Below we discuss the implementation details and the approximation of the diffusive model used to predict the variations of the humidity gradient in our experimental setup as shown in figures 4A and 4B of the manuscript. This model relies on the hypothesis that humidity transport uniquely happens by diffusion and happens in a quasi-stationary way, which correspond to solve the Laplace equation $\Delta h = 0$ for the humidity field $h$, which is done using the finite elements platform COMSOL Multiphysics. Our simulation domain was a cubic domain of side 18 mm whose bottom face is replaced by a 3D copy of the experimental setup, in the case of pillars cue (Fig. 4A) and wall cue (Fig. 4B). The boundary conditions on the field $h$ were no-flux on the lateral boundaries and Dirichlet on the bottom and top plate where the relative humidity $h$ was fixed to 100% and 70%, to mimic the experimental value of $h$ respectively at the clay disk surface and in the air (far from the disk surface) of our experimental room.

**Approximation of the diffusive zone** Note that the size of our simulation box overestimate by 10 times the thickness of the viscous boundary layer given in S11.

Our choice was made because there is no scale separation between our topographic cues and the boundary layer thickness and drawing the shape of the top boundary of the diffusive region is a very difficult task. Consistently, the value of $|\nabla h|$ on the disk surface is 10 times smaller of the estimation $|\nabla h|_0$ of the same quantity in our experiments reported in the manuscript. Note also, that as in all diffusive problems, the humidity gradient on any point of the bottom boundary (i.e. on the clay surface) depends on the distance of that point from the top boundary and the topography. Thus, in principle, the size of the simulation box does not only affect the overall magnitude of the humidity gradient but also its shape. However, one observes that in our simulations the topographic cues are only 30% closer to the top boundary compared to the flat, bottom, surface, but the local gradient is 10 to 20 times larger. This evidence suggests that the 'curvature' effect is much more important than the 'distance' effect, and supports the fact that our approximation does not affect in a significant way the estimation of the relative importance of the humidity gradient at the bottom surface. Conversely, the approximation clearly affects the absolute value of the humidity gradient, but this can be easily 're-scaled' using the reference value $|\nabla h|_0$.

### S.VII Comparison of simulations and data

Figure 2C reports the values of termite deposition activity from one experiment (red line, from experiment E66) alongside with the local surface curvature measured for the same experimental setup (blue line). The same figure panel also reports the deposition activity from a simulation of our 3D curvature-based growth model, initialized with the same topography as in experiment E66 (blue line). All values are plotted along a one-dimensional cross section cutting across the pillars.

The data contributing to this figure panel originate from different sources: experimental pellet collection and deposition events are obtained from time lapse photographs, while curvature is obtained from 3D scans. Below we explain how we realign the data to take the 1D projection shown in the figure. For the experimental pellet depositions, we simply considered the value of the deposition heatmap shown in figure 2A by selecting the cells adjacent to the black dashed line which is shown in the same figure. The value are then normalized by their maximum. For the curvature values, we took the values of curvature corresponding to the mesh elements shown in figure 3C which fall close to the same line. In order to quantify pellet depositions from the 3D simulations we proceed as follows: first, we identify all the voxels of added material by subtracting the original 3D shape from the final one. Then, we select all the points that fall close to the plane passing through the black dashed line shown in Fig. 2A. For each of these voxels, we then identify a *corresponding* voxel on the original 3D shape which is the closest one. Finally, we compute the radial positions in the horizontal plane of all the *corresponding* voxels and we plot the histogram of these radial values, where the histogram is normalized by its maximum to obtain the 1D projection shown in figure 2C.

### S.VIII Relationship between local surface curvature and evaporation field

Our chemical garden experiments, alongside with the stationary humidity field simulations described above in section S.VI both illustrate a relationship between substrate curvature and evaporation. In the following sections, we provide theoretical arguments in support for the relation between evaporation and curvature. In section S.VIII.A, we re-derive the expression for the humidity gradient at the surface of isolated spherical droplets of a given radius. Then, in section S.IX, we provide arguments demonstrating that similar results also apply with good approximation to our experimental substrate made available to termites.

#### S.VIII.A Diffusive evaporation from a spherical surface with a given curvature

As stated in the manuscript, the relationship between surface curvature and evaporation flux was observed by Langmuir [*S9*] already more than a century ago. Langmuir studied isolated droplets of an evaporating fluid and showed that the evaporation flux from an isolated droplet of radius $R$ is proportional to $R$, while the humidity gradient scales as $1/R$ at the droplet surface. For the sake of clarity this result is re-derived below following [*S7*]. Let $h(r)$ be the humidity field around the droplet, at the equilibrium the diffusion equation in spherical coordinates reads:

$$0 = \frac{1}{r^2}\frac{\partial}{\partial r}\left(r^2\frac{\partial h}{\partial r}\right) + \frac{1}{r^2\sin\theta}\frac{\partial}{\partial\theta}\left(\sin\theta\frac{\partial h}{\partial\theta}\right) + \frac{1}{r^2\sin\phi^2}\frac{\partial^2 h}{\partial\phi^2}. \tag{S1}$$

As the system has spherical symmetry, $\partial_\theta = 0$ and $\partial_\phi = 0$ thus diffusion equation reduces to:

$$0 = \frac{\partial}{\partial r}\left(r^2\frac{\partial h}{\partial r}\right), \tag{S2}$$

which means that humidity gradient $\partial h/\partial_r$ is $\partial h/\partial_r = B/r^2$ where $B$ is a constant, and the humidity field must have the form:

$$h(r) = A - \frac{B}{r} \tag{S3}$$

where $A$ is another constant. We assume then, that diffusion acts in a spherical shell between an inner sphere of radius equal to the radius of the droplet, and an outer sphere of radius $R_{out}$. Let $h(R) = h_0$ and $h(R_{out}) = h_{out}$; the constants $A$ and $B$ can then be obtained by substituting in S3 and they read:

$$A = h_0 - \frac{(h_0 - h_{out})R_{out}}{R_{out} - R} \quad B = \frac{(h_{out} - h_0)RR_{out}}{R_{out} - R}. \tag{S4}$$

We can then compute the humidity gradient at the droplet surface as:

$$\left.\frac{\partial h}{\partial r}\right|_R = \frac{(h_{out} - h_0)R_{out}}{R(R_{out} - R)}, \tag{S5}$$

now if we assume that $R_{out} \gg R$ one sees that the humidity gradient at the droplet surface scales like $1/R$ and that the humidity flux evaporating from the droplet must scale like $R$.

### S.IX Scaling arguments for a quasi-stationary diffusive regime

Our experimental system is more complex than isolated droplets because the evaporating substrate is a porous medium, and its geometry is more convoluted than an isolated sphere. However, the same conclusions are valid if two important conditions are fulfilled which are:

1 We restrict our attention to the viscous boundary layer;

2 The porous medium is equally soaked everywhere.

As we showed in figure 5, evaporation implies both humidity and temperature variations which can cause the onset of the so-called 'moist' convection, which is a complex phenomenon still under study by fluid-dynamicists [*S15*]. However, within the viscous boundary layer the system is dominated by diffusion. Below (subsection S.IX.A), we show that the thickness of the viscous boundary layer is comparable to the thickness of termites, so fulfilling condition 1. Condition 2 is also fulfilled. In fact, by combining Washburn's results [*S17*] for the dynamics of capillary rise in a tube and an effective value for the size of pores in our clay disk [*S5*], we can establish that there is a strong time scale separation between capillary rise through the clay disk and evaporation from the disk surface: evaporation is not limited by the capillary soaking of the substrate (section S.IX.B).

#### S.IX.A Estimation of the thickness of the viscous boundary layer

Previous experiments by Soar et al. [*S12*] indicated that termites live in the viscous boundary layer, that is, the thickness of the boundary layer is comparable with the typical body thickness of termites. Below we show that this assumption is justified also for our experiments with scaling arguments.

Traditionally, the thickness $\delta$ of the viscous boundary layer over a plate is estimated [*S2*] as:

$$\delta \propto \sqrt{\frac{l\nu}{v}}, \tag{S6}$$

where $l$ and $v$ are typical scales for the size of the system and the speed of the fluid and $\nu$ is the kinematic viscosity of the fluid. This scaling was successfully used to model water evaporation flux from a vessel [*S8*]. At 25 °C viscosity does not change much with relative humidity so we can take the value for dry air which is $\nu \sim 1.5 \cdot 10^{-5}\,\mathrm{m^2 s^{-1}}$ and we consider $l \sim 5 \cdot 10^{-2}\,\mathrm{m}$ which is the diameter of the clay disk. Estimating the velocity scale $v$ is a less easy task because, unlike in previous experiments by other authors [*S8*, *S12*], we are not imposing any air flow velocity in our experiments. However, this does not mean that the velocity is zero. In fact, above the viscous boundary layer humid, lighter air will flow upward pushed by the buoyancy force forming an uprising convective plume. By mass conservation, fresh dry air must come from the side of the disk with velocity which is parallel to the horizontal and comparable in magnitude with the velocity of the uprising plume. Air density is also affected by temperature, and as we showed in figure 5 there is a temperature drop at the clay disk surface because of evaporation which must increase air density and thus slow down or inhibit convection. For the sake of simplicity, we neglect temperature effects at this stage (though we will discuss them at the end of this section) which will give us an upper bound for $v$ and a lower bound for $\delta$.

To assess the buoyancy force we must quantify the density of humid air $\rho_d$. This can be estimated treating it as a mixture of perfect gases (Dalton's law) which leads to:

$$\rho = \rho_d - P_{sat}\phi \frac{R_d - R_v}{R_d R_v T}, \tag{S7}$$

where subscripts $d$ and $v$ indicate dry air and water vapor respectively, $R$ is the specific gas constant, $T$ is the absolute temperature, $\Phi$ is the relative humidity and $P_{sat}$ is the vapor saturation pressure. Using Teten's formula for computing $P_{sat} \sim 3.2 \cdot 10^3\,\mathrm{Pa}$ at 25°C, the density variation $\Delta\rho_v$ between the surface of the clay disk ($\Phi = 100\%$) and far from it ($\Phi \sim 70\%$) reads:

$$\Delta\rho_v \sim 4.4 \cdot 10^{-3}\,\mathrm{kgm}^{-3}. \tag{S8}$$

Using this value, we can estimate the Rayleigh number for our setup, which is:

$$Ra = \frac{g\delta_v H^3}{D_v \eta_a} \sim 3 \cdot 10^6, \tag{S9}$$

where $D_v$ is the mass diffusivity of water vapor in air, $\eta_a$ is the dynamic viscosity of air and the length $H$ is the height of the column of air between our experimental setup and the ceiling of the room. One observes that $Ra$ is much larger than the critical Rayleigh number for convection $Ra_c \sim 10^3$, thus if temperature effects are neglected, air is certainly convecting. We can then estimate the velocity induced by buoyancy copying the choice of [*S11*] for thermal convection:

$$v \sim \sqrt{gH\Delta\rho_v/\rho_d} \sim 0.25\,\mathrm{ms}^{-1}, \tag{S10}$$

which substituting in equation S6 gives

$$\delta \sim 2\,\mathrm{mm}, \tag{S11}$$

which is comparable with the body thickness of our termites ($\sim 1\,\mathrm{mm}$).

**Temperature effects.** Below we discuss the effect of evaporation cooling and we estimate the density difference caused by the temperature drop at the clay disk surface. According to figure 5, the temperature on the clay disk can be estimated to drop by about 1°C, while the expansion coefficient of air at 25°C is $\alpha \sim 3.4 \cdot 10^{-3}\,\mathrm{kgm}^{-3}$ thus the density difference induced by the temperature drop is

$$\Delta\rho_T = \rho_d \alpha \Delta T \sim 4.4 \cdot 10^{-3}\,\mathrm{kgm}^{-3},$$

which is very close to the value we find for the density difference induced by humidity (eq. S8). Note that this is a positive variation, i.e. cooler air is heavier, which should contrast and possibly inhibit convection. However, one should notice that if convection does not set, $\delta$ will be larger and the evaporation flux smaller as shown by equation S14. This means that, even if the cooling caused by evaporation slows down or possibly stops the evaporation from time to time, the process might start again following a sort of intermittent sequence. In any case, temperature effects can only weaken convection and thus increase the thickness of the the viscous boundary layer $\delta$, in which case the conclusions of our paper are nothing but reinforced.

#### S.IX.B Capillary rise through a porous medium

In the manuscript we stated that the clay disk is constantly replenished in water by capillary rise while evaporating from the surface. In particular, we assume that this is a quasi stationary process and that the value of moisture at the disk surface is approximately constant and air is saturated with vapor. This assumption relies on a strong scale separation between the typical time of evaporation and the time that water needs to rise up to the disk surface. Below we show that such a scale separation does exist in our experiments.

It is well known that a fluid can flow through a capillary up to a equilibrium height $H_c$ which is given by the following formula:

$$H_c = \frac{2\gamma \cos\theta}{\rho g r} \tag{S12}$$

with $\gamma$ the surface tension, $\theta$ the contact angle, $\rho$ the fluid density, $g$ the gravity and $r$ the capillary diameter. The case of a porous medium is more complex but equation S12 can still be used choosing the adequate *effective* pore radius $r$, and we take $r \approx 10^{-7}\,\mathrm{m}$ [*S5*]. Clay is hydrophilic, which means that we can approximate $\cos\theta \approx 1$ and we remind that surface tension for water at 25 °C is $\gamma \approx 7{\cdot}10^{-2}\,\mathrm{Nm^{-1}s^{-1}}$ and $\rho g \approx 10^4\,\mathrm{Nm^{-3}}$. One can then estimate $H_c$ as $H_c \approx 100\,\mathrm{m}$ in clay. One will notice that in our experimental setup water must climb no more than 6 mm from the reservoir to the top of our topographic cues thus the mechanism of capillary rise appear more than effective to constantly replenish the clay disk with water and gravity can be neglected in the rising dynamics. With such hypothesis Washburn [*S17*] showed that the capillary rise $h$ evolves with a diffusion law $h^2 = D_\gamma t$ where the effective diffusion coefficient $D_\gamma$ can be written as:

$$D_\gamma = \frac{\gamma \cos\theta r}{2\eta_w} \approx 4 \cdot 10^{-6}\,\mathrm{m^2s^{-1}}, \tag{S13}$$

where $\eta_w = 10^{-3}\,\mathrm{Nm^2s^{-1}}$ is the dynamic viscosity of water. As we stated in the manuscript, close to the surface the uptake of water also happens by diffusion. The time scale separation between capillary rise and evaporation can then be estimated computing the corresponding diffusive flux $q_\gamma$ and $q_{ev}$ across the clay disk thickness $d$ and the viscous boundary layer $\delta$ respectively:

$$q_\gamma = \frac{SD_\gamma \rho_l}{d}, \qquad q_{ev} = \frac{SD_v \Delta\rho_v}{\delta}. \tag{S14}$$

Approximating the surface $S$ to be the same for the two processes and taking $d = 6\,\mathrm{mm}$ as an upper boundary for the clay disk thickness (i.e. the height of the topographic cues), the ratio between the two fluxes reads:

$$\frac{q_\gamma}{q_{ev}} \sim \frac{D_v}{D_\gamma}\frac{\Delta\rho_v}{\rho_l}\frac{1}{3} \sim 10^3. \tag{S15}$$

where we use the value of $\delta$ and $\Delta\rho_v$ obtained in section S.IX.A, the effective diffusion $D_\gamma$ from equation S13 and $D_v = 2.6 \cdot 10^{-5}\,\mathrm{ms}^{-2}$ the mass diffusivity of vapor at 25°C [*S4*]. One observes that time scales of capillary rise and evaporation are separated by three order of magnitude which means that our assumption is highly reliable. As an example it would remain valid even if we were about to commit an error of 2 order of magnitude in (over) estimating $r$ the effective pore size of our clay, which is by far the highest source of uncertainty in this estimation. Finally note that the time scale of capillary rise is a decreasing function of the pore size $r$ (see eq. S13), while $H_C$ scales like $1/r$ (see eq. S12), thus our assumptions remain valid even if the new construction made by termites included bigger pores up to a size of 1 mm.

[S2] D. J. Acheson. Elementary Fluid Dynamics. Oxford University Press, Oxford : New York, 1990.

[S3] D. S. Calovi, P. Bardunias, N. Carey, J. Scott Turner, R. Nagpal, and J. Werfel. Surface curvature guides early construction activity in mound-building termites. Philosophical Transactions of the Royal Society B: Biological Sciences, 374(1774):20180374, 2019.

[S4] E. L. Cussler. Diffusion: Mass Transfer in Fluid Systems. Cambridge University Press, 1997.

[S5] S. Diamond. Pore Size Distributions in Clays. Clays and Clay Minerals, 18(1): 7–23, 1970.

[S6] G. Facchini, A. Lazarescu, A. Perna, and S. Douady. A growth model driven by curvature reproduces geometric features of arboreal termite nests. Journal of The Royal Society Interface, 17(168):20200093, 2020.

[S7] N. Fuchs. Concerning the Velocity of Evaporation of Small Droplets in a Gas Atmosphere. Technical Report NACA-TM-1160, 1947.

[S8] K. Hisatake, S. Tanaka, and Y. Aizawa. Evaporation rate of water in a vessel. Journal of Applied Physics, 73(11):7395–7401, 1993.

[S9] I. Langmuir. The Evaporation of Small Spheres. Physical Review, 12(5):368–370, 1918.

[S10] H.-F. Li, N.-Y. Su, and W.-J. Wu. Solving the hundred-year controversy of coptotermes taxonomy in taiwan. American Entomologist, 56(4):222, 2010.

[S11] W. V. R. Malkus and M. C. Thayer. Notes on the 1964 Summer Study Program in Geophysical Fluid Dynamics at the Woods Hole Oceanographic Institution. Technical Report, Woods Hole Oceanographic Institution, 1964.

[S12] R. Soar, G. Amador, P. Bardunias, and J. S. Turner. Moisture gradients form a vapor cycle within the viscous boundary layer as an organizing principle to worker termites. Insectes Sociaux, 66(2):193–209, 2019.

[S13] N.-Y. Su, P. Ban, and R. H. Scheffrahn. Resistance of insecticide-treated foam board insulation against the eastern subterranean termite and the formosan subterranean termite (isoptera: Rhinotermitidae). Journal of Economic Entomology, 96(5):1526–1529, 2003.

[S14] N.-Y. Su, T. Chouvenc, and H.-F. Li. Potential hybridization between two invasive termite species, coptotermes formosanus and c. gestroi (isoptera: Rhinotermitidae), and its biological and economic implications. Insects, 8(1):14, 2017.

[S15] G. K. Vallis, D. J. Parker, and S. M. Tobias. A simple system for moist convection: The Rainy–Bénard model. Journal of Fluid Mechanics, 862:162–199, 2019.

[S16] T. Walter and I. D. Couzin. TRex, a fast multi-animal tracking system with markerless identification, and 2D estimation of posture and visual fields. eLife, 10:e64000, 2021.

[S17] E. W. Washburn. The Dynamics of Capillary Flow. Physical Review, 17(3): 273–283, 1921.
